# A Social Affordance Framework for Understanding Approach Decisions in Wild Crested Macaques (*Macaca nigra*)

**DOI:** 10.64898/2026.06.25.734458

**Authors:** Adam Provin, Fannie Beurrier, Muhamad Julfikram Bawimbang, Dyah Perwitasari-Farajallah, Sébastien Ballesta, Cécile Garcia, Julie Duboscq

**Affiliations:** UMR 7206 Eco-Anthropology Centre National de la Recherche Scientifique, Muséum national d’Histoire naturelle, Université Paris Cité – Musée de l’Homme - 17 place du Trocadéro - 75016 Paris, France; Macaca Nigra Project Tangkoko Reserve - Bitung - 95535 - North Sulawesi - Indonesia; Faculty of Mathematics and Natural Sciences Biology Department - IPB University – Jl. Raya Dramaga, Kampus IPB Dramaga Bogor, 16680 West Java, Indonesia; Primate Research Centre IPB University – Jl. Lodaya II/5, Bogor 16151, Indonesia; Centre de Primatologie de l’Université de Strasbourg Chemin du Fort Foch - 67207 - Niederhausbergen, France; UMR 7364 Laboratoire de neurosciences cognitives et adaptatives Université de Strasbourg, Centre National de la Recherche Scientifique Faculté de psychologie 12 rue Goethe - 67000 - Strasbourg, France

**Keywords:** approach decisions, social affordances, partner choice, macaques

## Abstract

Approaching conspecifics is a fundamental component of social interactions. Such decisions are typically investigated through the study of social relationships (e.g., kinship, dominance, affiliation) and their impact. In the framework, the role of social affordances, i.e., interaction opportunities arising from the immediate social environment, remains poorly understood. We investigated how context-dependent factors shape approach decisions in wild crested macaques (*Macaca nigra*) in North Sulawesi, Indonesia. From September 2024 to May 2025, we collected data on 27 adults (20 females, 7 males) using focal observations and an approach-specific protocol recording neighbours’ identities and behaviours at the initiation of close-proximity approaches (<1 body length). We first compared data collected during approaches with focal data, then conducted within-approach analyses contrasting approached partners and bystanders. We further tested the effects of behavioural state, subgroup size and composition, including infant presence. Bayesian models revealed that affiliative, neutral, and grooming behaviours were more frequent in approach contexts than in general observations. Within approaches, individuals avoided congeners already engaged in grooming and approached individuals in neutral or affiliative states (e.g., lip-smacking, embracing). Macaques globally approached more isolated individuals or dyads. When multiple subgroups were present (i.e., multiple choices), the likelihood of being approached increased with subgroup size and was higher in subgroups containing infants. These findings indicate that macaques use behavioural cues and local social density as affordance signals, adjusting approach decisions to maximize interaction opportunities and navigate complex social environments.

## Introduction

Many non-human primate species live in stable social groups, enabling the formation of preferential social bonds, which are positive relationships between individuals through repeated interactions with selected conspecifics. These social bonds are often associated with increased fitness, resource sharing, and mating opportunities (Silk et al., 2013; Silk, 2007). These bonds are built through repeated partner choices that can be driven by multiple factors, including kinship, relationship strength, individual preferences, partners’ skills or services, mating opportunities, and identity traits such as age or sex (Mielke et al., 2018; Molesti & Majolo, 2016; Siracusa et al., 2022; Suchak et al., 2014). Approaching conspecifics is a prerequisite for social interaction and can be considered as a social decision that brings individuals close enough to enable most interactions (Farine, 2015), i.e., a partner choice. An approach is thus the movement that brings one individual into close proximity to another (i.e., within one body length), thereby increasing the likelihood of visual, auditory, or tactile contact and facilitating subsequent social interaction. In primates, these approach dynamics were first studied in mother-infant interactions, recording the number of times the infant and the mother approached or moved away from each other (Hinde et al., 1964), and have then been extended to other individuals in the group (Hinde, 1977). While the importance of social relationship characteristics (kinship, dominance difference, affiliation strength) in shaping partner choice is well established, approach dynamics can also be driven by contextual factors that may influence who can be approached at any given moment, such as local social density (e.g., the number of other individuals in vicinity), the composition of nearby groups (e.g., presence or absence of an infant), or the current behavioural states of the individuals composing these groups (e.g., foraging versus grooming). These contextual constraints highlight the concept of social affordances, defined as the interaction opportunities arising from the social environment, as perceived by the individual. Originally introduced by Gibson as the action possibilities offered by inanimate objects to organisms, the affordance concept was subsequently extended to encompass opportunities provided by animate agents, including conspecifics (Gibson, 1979; Orban et al., 2021; Schilbach et al., 2013). Aforementioned affordances can be direct signals, such as facial expressions, inviting to enter into interaction. For instance, in wild chimpanzees (*Pan troglodytes verus)* and mangabeys (*Cercocebus atys atys*), individuals moving within close range exchange signals (e.g., visual signals, such as head movements, present body ; or auditory, like bark hoo, pant-grunts in chimpanzees and twitter or growl in mangabeys) that help to negotiate tolerance and cooperation, and the reciprocity of such signalling covaries with relationship strength (Grampp et al., 2025). These facial expressions can convey positive or aggressive intentions or playful intensities (Bresciani et al., 2022). However, facial expressions can also have different meanings depending on the context: silent-bared teeth in rhesus macaques (*Macaca mulatta*) are perceived as signals of subordination that reduce the likelihood of direct aggression and promote grooming in peaceful contexts, but function as immediate submission during agonistic confrontations (Beisner & McCowan, 2014). Social affordances can also take the form of body postures, known as bodily affordances (Van Boekholt et al., 2024). They can promote or facilitate an action: chimpanzees play coordination emerges when one individual’s postural alignment (e.g., nesting, hanging) invites reciprocal moves, co-constructing joint activity without discrete signals (Van Boekholt et al., 2024). In some cases, individuals can develop specific behaviours to indicate their availability or non-availability: isolated individuals from a mandrill community (*Mandrillus sphinx*) have been observed covering their eyes with their hands, with the consequence to be directly less touched and approached by others (Laidre, 2011). Monkeys may then perceive some behaviours as social affordances that indicate whether a partner is available to interact at a given moment. Finally, social affordances may themselves be modulated by environmental or social contexts, such as group composition or local social density, which could shape the action opportunities available to individuals. One of the most studied factors related to social opportunities or the likelihood of interaction is infant presence (Dunayer & Berman, 2018). Indeed, infants can alter partner-choice dynamics and approach motivation, by attracting attention, or influencing partner availability (Maestripieri, 1994; Pfaff et al., 2025). For example, female Tibetan macaques (*Macaca thibetana*) with infants exhibit higher grooming rates than those without infants (Jiang et al., 2019). Within a biological market framework (Noë & Hammerstein, 1994), infants attract interest from others seeking handling or bonding opportunities, leading non-mothers to direct more grooming toward mothers and increasing the mother’s social value within the group. From a social affordance perspective, the presence of an infant gives opportunities for infant handling or playing, and potentially, relationship strengthening with nearby adults and/or the infant’s mother. In contrast to a simple increase in social value, infants create new affordances by turning mothers into distinct social opportunities, which in turn reshapes the group’s interaction landscape and increases the probability of being approached. Besides infant presence, local neighbour densities are a key determinant of interaction opportunities, but most work has focused on spacing, grooming networks and predator vigilance rather than on approach rates. In wild Assamese macaques (*Macaca assamensis*), individuals had fewer close neighbours and larger nearest-neighbour distances when the group was feeding than when resting or moving, indicating that animals actively reduce local density in a food competition context (Heesen et al., 2015). Juveniles and females with young infants, who are more vulnerable to predation, were found located closer to the centre of the group and surrounded by more neighbours than adult males and females without infants, showing that vulnerable individuals seek higher local densities to reduce predation risk (Heesen et al., 2015). Similar anti-predator strategies occur in crested macaques *(Macaca nigra)*, where individuals collectively gather and mob a reticulated python (*Malayopython reticulatus*) after hearing alarm calls from conspecifics (Micheletta et al., 2012). In baboons and other primates, age, sex and dominance shape consistent spatial positions and neighbourhood sizes, with more central individuals interacting with a larger number of neighbours (Farine et al., 2017; Pereira, 1988). Relatively few studies have examined how the local social density around potential partners affects an individual’s decision to initiate social approaches. Analysing approach decisions as a function of both the audience and the potential partner’s local social density would shed light on how individuals chose their social partners and structure their social group.

Together, these previous studies indicate that immediate behavioural, environmental, and spatial contexts strongly modulate when and how individuals can initiate contact, either for affiliative interactions (e.g., grooming) or for agonistic ones (e.g., aggression). However, despite their importance, these immediate factors remain understudied relative to kinship and relationship characteristics in partner-choice research. In this study, we investigated factors influencing approach behaviour and, consequently, partner choice, in a tolerant macaque species, the crested macaque (*Macaca nigra*). The macaque genus is particularly interesting for investigating these immediate factors, as they live in large, hierarchical, matrilineal groups with diverse levels of social tolerance that could influence the way they choose partners. Macaques are classified in a 4-grade scale, from intolerant (Grade 1: high intensity conflicts, low/no conciliatory tendencies, strong dominance asymmetry, strong nepotism; e.g., *Macaca fuscata*) to tolerant (Grade 4: low intensity conflicts, high conciliatory tendencies, moderate dominance asymmetry, weak nepotism; e.g., *Macaca nigra*) (Thierry, 2007, 2015). This gradient creates rich opportunities to study approach behaviours and social affordances: as tolerant species allow close proximity to diverse partners, they could show context-specific partner choice beyond kinship and dominance, in contrast to more intolerant species where individuals are more constrained in their partner choice freedom. Moreover, infant attractiveness varies across this gradient, with a strong influence of kinship and rank on infant handling opportunities in intolerant species, but with more diverse handlers in tolerant macaques (Dunayer & Berman, 2018). Crested macaques offer an ideal system for this question because tolerant species are expected to have more flexible social interactions and broader access to potential partners. This increases the likelihood of observing how social affordances influences approach decisions beyond kinship and dominance.

To investigate social affordances, we examined whether potential partners’ current behavioural state, audience size (number of neighbours), and the presence of infants influenced approachers’ decisions. By doing this, we tried to determine which factors are perceived as social affordances by crested macaques. First, we compared the occurrence of behaviours during baseline focal observations with the occurrence of those observed at the moment of an approach, expecting that if individuals preferentially approach others in specific behavioural states, behavioural profiles during approach would differ from those recorded during baseline observations. We then tested whether chosen partners’ behavioural states differed from those of surrounding audience members, with the prediction that affiliative displays and neutral states would be more frequent in partners. Second, to examine the influence of neighbours’ presence, i.e., local density, we compared neighbour counts around focal individuals between baseline and approach contexts, predicting that chosen partners would have fewer neighbours than audience individuals, if we consider that high local density offers less possibilities of interactions than a lower one. Finally, we investigated how subgroup composition influenced approaches and tested whether approach patterns varied with subgroup size and infant presence. To distinguish infant-specific effects from broader subgroup size effects, and given that larger subgroups are more likely to include infants, we repeated this analysis using only adult approaches (no infants present in either partner or audience).

## Material and Methods

### Ethical Statement

This research adheres to all legal requirements and guidelines of the French and Indonesian governments (Indonesian National Research and Innovation Agency, Permit Number 305/SIP/IV/FR/5/2024, Ethical Clearance Approval Number 077/KE.02/SK/03/2024).

### Field site and study population

A wild population of crested macaques (*Macaca nigra*) has been studied in the Tangkoko Nature Reserve, North Sulawesi, Indonesia (1°33’N, 125°10’E), within the Macaca Nigra Project (Duboscq & Micheletta, 2023), a long-term field project focusing on the biology, ecology and conservation of crested macaques in Tangkoko since 2006. These macaques live in multimale-multifemale groups of around 20 to 100 individuals, with philopatric females organised in stable dominance hierarchies (Duboscq et al., 2017). They inhabit lowland rainforest characterised by a highly seasonal climate and fluctuating food availability, and spend around 60% of their time on the ground (Joly et al., 2017; O’Brien & Kinnaird, 1997). Crested macaques are classified as Critically Endangered on the IUCN Red List of endangered species (IUCN, 2020). For our study, we followed one group (PP) from sleeping site to sleeping site, from dawn to dusk, five days per week. The group comprised approximately 50 individuals, including 20 adult females, 7 adult males, and around 20 juveniles and infants. The individuals are well habituated to human observers and are identified individually based on their physical characteristics alone. They were never approached closer than two meters and observers did not interact with the animals nor did they interfere with the expression of their natural behavioural repertoire.

### Observational data collection

We observed all individually recognisable adults in the group. While one observer (*Anonym 1*) was collecting behavioural data using focal sampling (Altmann, 1974), a field assistant (*Anonym 2*) collected data during approach events using a protocol specifically designed for this study. Focal observations consisted of 30-minute point-sample observations of a single individual. During each observation, we recorded the subject’s activity at one-minute point intervals and continuously recorded all occurrences of social and self-directed events (Duboscq et al., 2013, 2017). Activities included foraging (searching for food), food processing (manipulating food), feeding (ingesting food and chewing), drinking (swallowing water), travelling (walking, running, jumping, or climbing), socialising (positive social behaviour, such as affiliative behaviour, grooming, playing), sexual behaviours (jaw-movement, sexual presentation, mating), aggressive interaction (threats, hits, chases, support shakes), staying immobile or resting (standing, lying down or sitting without obvious other behaviours) and self-grooming (self-inspection of the fur). Every minute, we also noted down the identity of neighbours according to three categories: in body contact, within one body length, and within five body lengths (Duboscq et al., 2013).

To record detailed data on approach events, we used a protocol specifically designed for this study, which recorded the identity of neighbours at each distance category and the behaviours (see definitions above) of the partner(s) and the audience (**Fig.1**). Each time an individual approached a conspecific at a distance of one body length, FB recorded the identity of the approacher and their behavioural state immediately after approaching, the identity of the partner, and all individuals composing the audience. The behaviours and positions of all individuals were noted using the same categories as in the basic focal sampling protocol: foraging, feeding, grooming, affiliative, aggressive, sexual, immobile or in motion (locomotion), self-directed behaviours (self-grooming or scratching), and we added infant- related behaviours (such as nipple suckling or holding). This resulted in a position matrix that provided information on both the distance and the behaviours of each individual at the precise moment of an approach.

**Figure 1:**
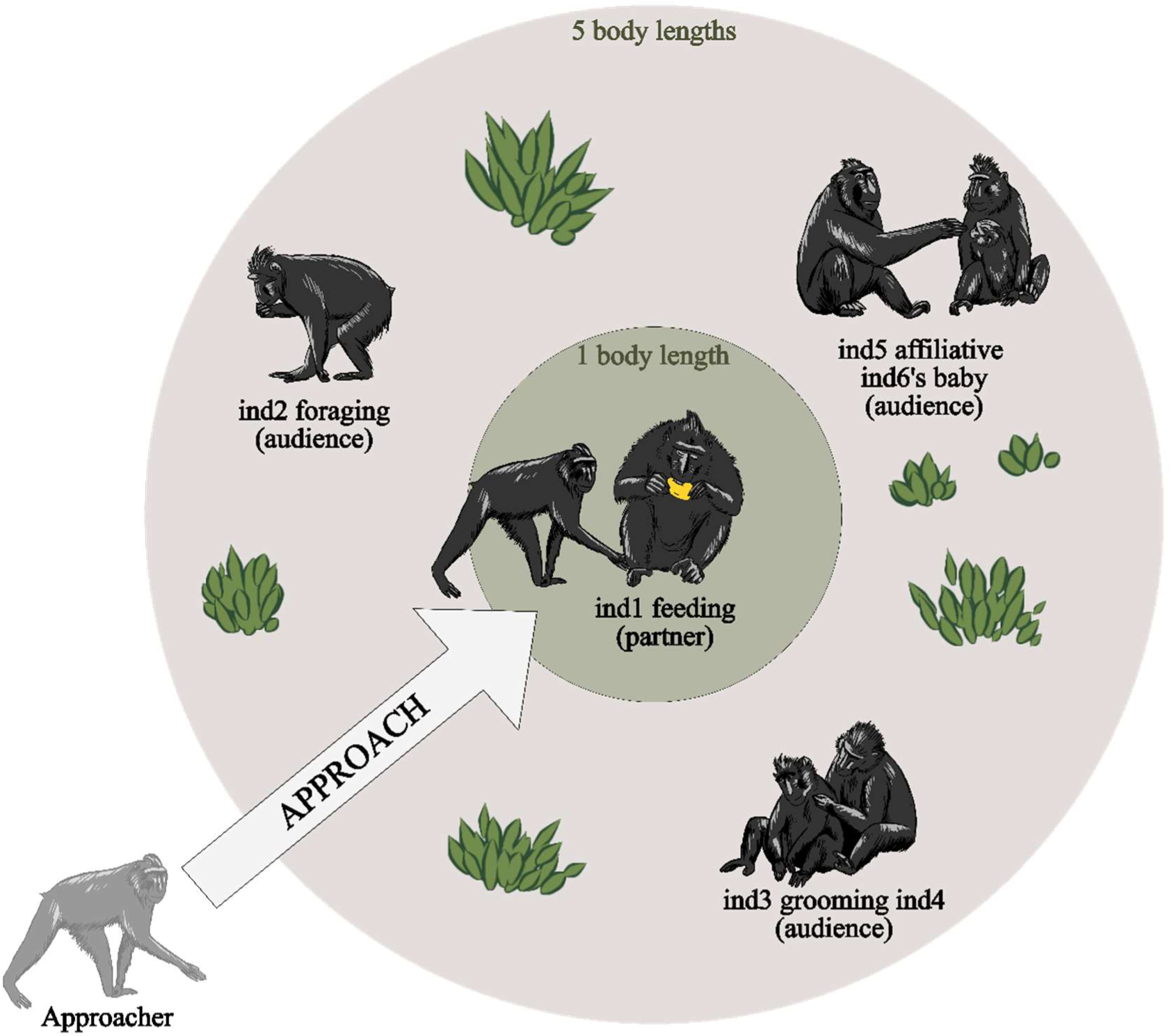
Approach protocol used during data collection on *Macaca nigra* at Tangkoko, Indonesia, January - May 2025. At the moment an individual (approacher) came in close proximity (within 1 body length) to a congener, we recorded the identity of the approacher, the behavioural state and identity of the partner (chosen individual, within the 1 body length range) and the audience (individuals in a five-body length range of the partner). In this example, individual “ind2” was five body lengths away from “ind1”, “ind3” and “ind4” were in body contact, “ind5” was within one body length of “ind6” and her baby, and “ind1”, “ind2”, “ind5”, “ind6” and her baby were at five body lengths. Macaques’ illustration by Mathilde Haeuser.

Focal data were collected from September 2024 to May 2025, and the approach protocol was implemented from January to May 2025, and conducted simultaneously with the focal protocol. 681 focal observations were collected on 27 adult individuals (356.67 hours in total; 12.74 ± 0.52 hours per individual). We recorded 1593 approaches (59 ± 6.44 per individual) with a total of approximately 45 different approachers (27 adults, that were followed also during the focal sampling, 14 identified infants, 2 identified juveniles, and other unidentified juveniles and infants).

### Statistical analyses

Analyses were conducted in R 4.5.2 (R Core Team, 2025) using RStudio 2025.09.1 (Posit team, 2025) using the packages ‘rstanarm’ (version 2.32.7) and ‘brms’ (version 2.23.0). For each analysis, we fitted a full model and a corresponding null model. The null model contained the same random effects and offset structure, but excluded the fixed effects. We compared full and null Bayesian models using leave-one-out cross-validation (LOO-CV; Vehtari, Gelman & Gabry, 2017), implemented in the loo package (version 2.8.0), and evaluated differences in expected log predictive density (ELPD) to assess whether the predictors of interest improved model performance beyond baseline variation. We used the conditional_effects function from ‘brms’ to visualise the fitted relationships and interaction patterns predicted by the Bayesian models, and the pairs function from ‘emmeans’ to obtain pairwise comparisons and assess whether specific contrasts between conditions were credibly different from zero. For this study, we deliberately did not account for social relationships and individual characteristics (e.g., sex, age), treating all monkeys as equivalent, to isolate the effects of social affordance factors “all else being equal”.

### Influence of behavioural states on approaches

To test whether behavioural states differ between approaches and baseline observations, we compared behaviour occurrences between these two datasets. We fitted a Bayesian zero-inflated negative binomial mixed model to account for overdispersion and zero inflation in the behaviour count data (Bekalo & Kebede, 2021; Rodríguez, 2022). The response was the occurrence of behaviours (counts) with two predictors, the context (binary: focal or approach), the category of the behavioural state (affiliative, aggressive, feeding, foraging, grooming, neutral and sexual) and their interaction. The counts were grouped by month and by individual, and these variables were included as random intercepts to account for potential monthly variation in behavioural states and repeated observations of individuals. We included the interaction between behavioural state and context to test our core prediction: if individuals preferentially approach congeners displaying specific behavioural states, certain categories will be overrepresented (e.g., affiliative interaction) or underrepresented (e.g., agonistic, non-social behaviours) during approaches compared to baseline. We also included an offset (log(Total)) to normalise observation effort between focal data (one line per minute) and approach data (only when an approach happened)(Winter & Bürkner, 2021). A second model focused on approaches only, comparing the behaviour of the partner with the behaviour of individuals in the audience, to test whether individuals preferentially approach congeners engaged in specific behaviours. The model structure was the same as the previous one, except that the predictor of interest was the individual’s role during the approach, coded as partner or audience. The interaction between the role and the behavioural state tests whether chosen partners show different behavioural states than audience members, particularly so for certain behavioural states compared to others. For example, the social affordance hypothesis predicts that partners will have higher rates of affiliative behaviours, whereas audience members may show higher rates of grooming (already engaged and therefore not being considered as available).

### Influence of social density on approaches

In order to test whether individuals preferred approaching denser subgroups, we investigated whether they had more social neighbours during approaches than during baseline focal sampling, and also between partner and audience. The first model tests whether approaches occur in denser contexts overall or target isolated individuals, while the second model tests whether chosen partners are relatively isolated compared to the nearby audience. To do so, we fitted a negative binomial model with the number of individuals in response, the context (focal or approach) and the distance category of neighbours in proximity (body contact, one body length, five body lengths) as predictors. We included a context × distance interaction to test whether approaches occur preferentially when more individuals are clustered at close proximity (body contact, one body length) than at five body lengths compared to baseline observations. We have two competing hypotheses that predict opposite patterns: 1. crested macaques being tolerant, approaches could occur in denser social contexts than during focal observations; or 2. Macaques approach isolated individuals to avoid interaction interruptions and competition for social interaction. As the previous models, we included month and individuals as random intercepts and an offset. For comparison, we fitted another model in which the context predictor was replaced by the individual’s role during the approach, distinguishing partner from audience. We hypothesized here that partners could have fewer individuals at close proximity, allowing for interaction.

### Influence of subgroup composition on approaches

To test the influence of subgroup composition (here specifically, number of individuals and the presence of an infant), we divided potential partners into subgroups as they were distributed spatially at the moment of the approach: a subgroup being constituted of individuals one body length apart from each other (can be also one individual without neighbours). We restricted analyses to approaches occurring when at least two subgroups were present, so that approachers had multiple choices, resulting in 533 approaches. We compared two Bayesian models: a model that predicted the binary probability of being approached as a function of subgroup size (number of individuals) in interaction with infant presence (binary: infant or no infant) to test whether infant presence amplifies or reduces the attractiveness of subgroups; and a model without interaction to test the effects independently. We also extracted adult-only approaches (both approacher and approached individuals) for a parallel analysis testing subgroup size effects independent of infant presence. We used a Bernouilly family (link logit) incorporating random intercepts for approacher identity and each approach event to take into account variation between approaches and approacher identity repeatability. In all analyses, models’ diagnostics were good (all R-hat = 1.00, ESS > 1000), and dispersion was controlled for each fitted model. Full results can be found in the Electronic Supplementary Materials.

## Results

### Influence of behavioural states on approaches

Model comparisons using leave-one-out cross-validation showed that the full model outperformed the null model in both analyses, indicating context-specific behavioural differences (ΔELPD = 1300.8, SE = 36.5) and role-specific behavioural states within approaches (ΔELPD = 394.1, SE = 17.2). (**Fig.2**; see the Electronic Supplementary Materials for full ELPD comparison tables).

**Figure 2.**
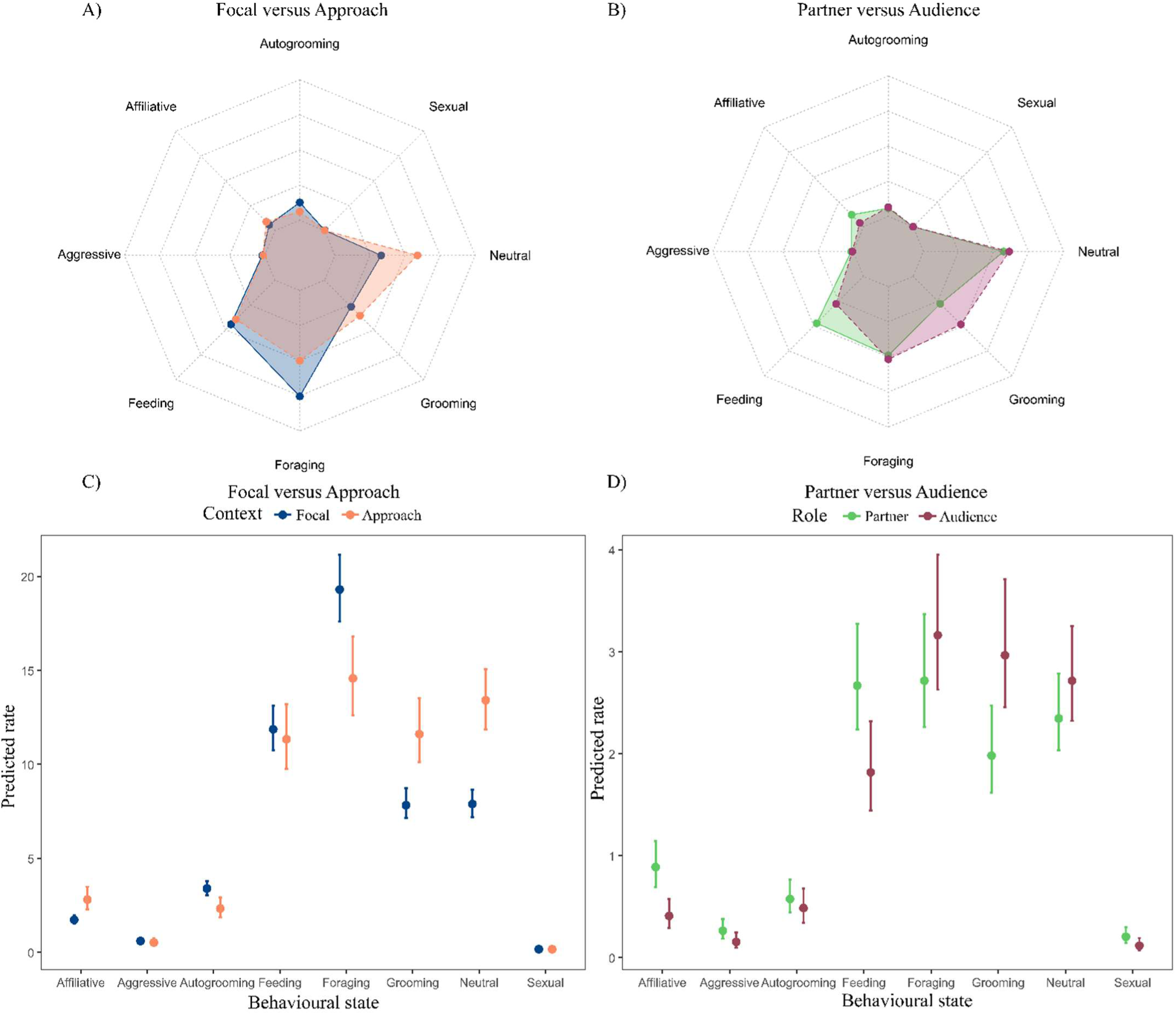
Proportions and conditional effects of behaviours across contexts. (Focal vs. Approach) and roles (Partner vs. Audience) in *Macaca nigra* at Tangkoko, Indonesia (September 2025 - May 2026). Radar plots show mean proportions of behaviours between focal and approach (**A**) and between partner and audience (**B**). Radar axes are scaled to a maximum of 0.5 for visual comparability across polygons. Conditional effects from the models are shown for focal versus approach (**C**) and partner versus audience (**D**). Axes show predicted neighbours per unit across distance categories. Points represent model-predicted rates, and error bars indicate 95% credible intervals.

### Focal versus approach

Radar plots and conditional effects highlights differences in behaviours between focal and approach contexts (**Fig.2**). Pairwise contrasts of estimated marginal means (EMM Focal - EMM Approach) revealed that only self-grooming (ΔELPD = 1.07, 95% HPD: [0.41, 1.68]) and foraging (4.69, 95% HPD: [2.13, 7.34]) were credibly higher in focal than approach observations. In contrast, neutral (-5.51, 95% HPD: [-7.05, -3.88]), affiliative behaviour (i.e., affiliative facial expressions, ΔELPD = -1.07, 95% HPD: [-1.68, -0.44]) and grooming (-3.76, 95% HPD: [-5.73, -2.11]) were more frequent during approaches. Feeding, aggressive and sexual behaviour did not differ between focal and approach contexts.

### Partner versus audience

Radar plots and conditional effects showed specific behaviours’ differences between partner and audience (**Fig.2**). Pairwise contrasts of estimated marginal means (EMM Partner - EMM Audience) showed that affiliative behaviour (ΔELPD = 0.47, 95% HPD: [0.24, 0.70]) and feeding (0.86, 95% HPD: [0.30, 1.44]) were credibly more frequent in partners than in the audience. Aggressive (0.11, 95% HPD: [0.02, 0.21]) and sexual behaviour (0.09, 95% HPD: [0.01, 0.17]) were also more frequent in partners than in the audience, but the effect was weaker. Grooming was less frequent in partners than in audience (-0.97, 95% HPD: [-1.16, 0.33]). Neutral behaviour, self-grooming and foraging did not differ between partners and audience individuals.

### Influence of social density on approaches

Model comparisons using leave-one-out cross-validation demonstrated that the full model outperformed the null model in both analyses. This provides evidence that neighbour density patterns differ systematically between baseline focal observations and approach contexts across distance categories (body contact, 1 body length, 5 body lengths) (ΔELPD = 321.8, SE = 24.8) and between roles across distance categories (ΔELPD = 461.1, SE = 31.6) (**Fig.3**; see the Electronic Supplementary Materials for full ELPD comparison tables). Mean number of neighbours per individual barplots can be found in the Supplementaries.

**Figure 3.**
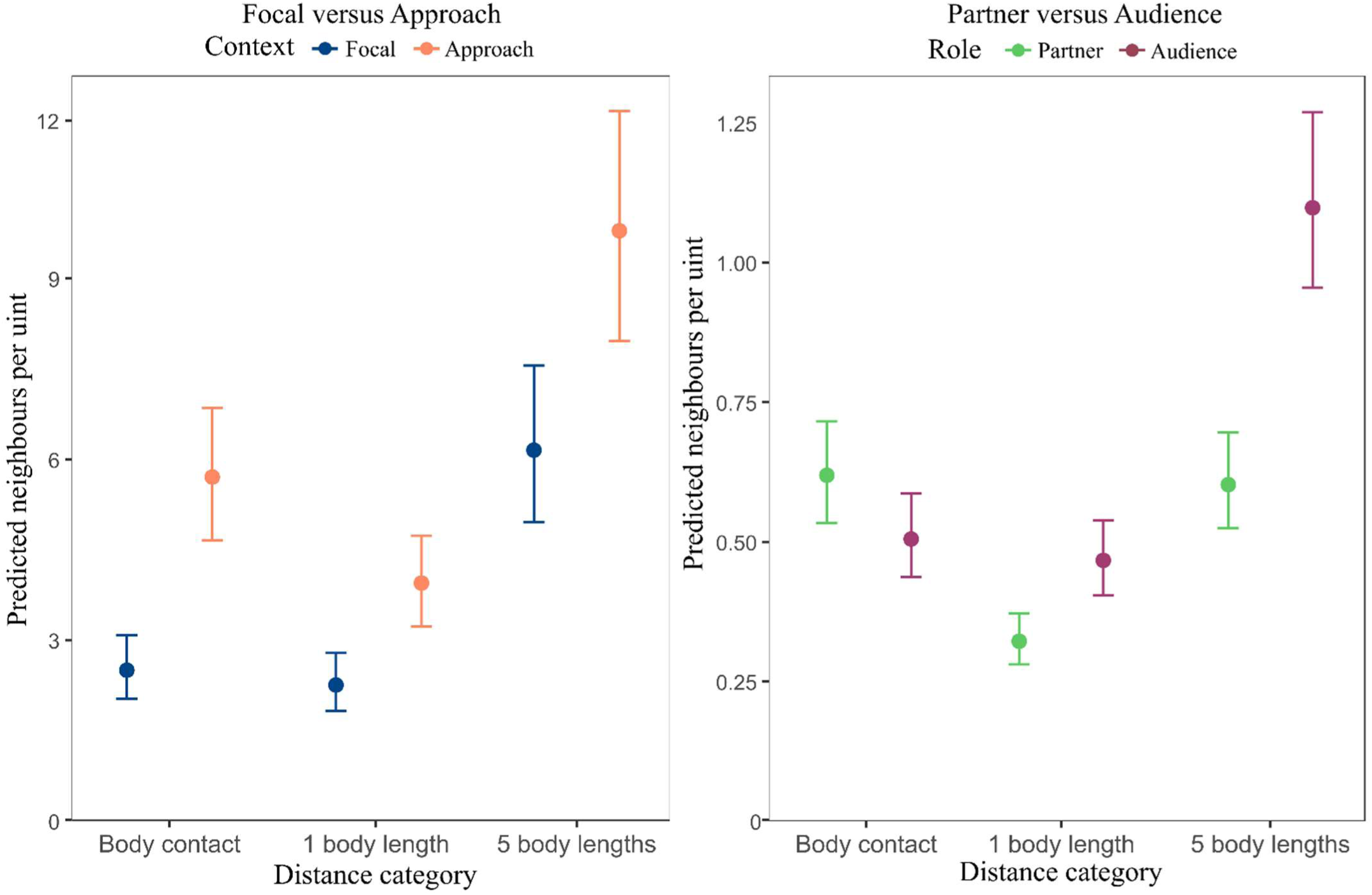
Conditional effects of neighbours per distance categories across contexts. (Focal vs. Approach) and roles (Partner vs. Audience) in *Macaca nigra* at Tangkoko, Indonesia (September 2025 - May 2026). Conditional effects from the models are shown for focal versus approach (**A**) and partner versus audience (**B**) Axes show predicted neighbours per unit across distance categories. Points represent model-predicted rates, and error bars indicate 95% credible intervals.

### Focal versus approach

Neighbour density was strongly associated with both context and neighbour category. Relative to the focal context, the approach was associated with higher neighbour counts overall (β = 0.82, 95% CI [0.66, 0.98]). Models showed moderate variation across individuals (σ = 0.35, 95% CI [0.25, 0.50]) and across months (σ = 0.14, 95% CI [0.05, 0.31]). Pairwise contrasts of estimated marginal means (EMM Focal - EMM Approach) showed that there were fewer neighbours during focal observations than in approach observations at all distance categories (body contact: ΔELPD = -3.19, 95% HPD: [-4.01, -2.41]; 1 body length: -1.69, 95% HPD: [- 2.27, -1.17]; 5 body lengths: -3.61, 95% HPD: [-4.97, -2.37]).

### Partners versus audience

Neighbour density depended on both social role and distance category. At the reference distance of body contact, audience individuals had fewer neighbours than partners (β = -0.20, 95% CI [-0.29, -0.12]). Moderate variation occurred across approach events (σ_app ≈ 0.57), whereas much smaller variation occurred across individuals (σ_ID ≈ 0.09) and months (σ_month ≈ 0.10). Pairwise contrasts of estimated marginal means (EMM Partner - EMM Audience) showed that partners had higher neighbour counts than audience members at body contact (ΔELPD = 0.11, 95% HPD [0.06, 0.16]), whereas they had lower neighbour counts than audience at 1 body length (ΔELPD = -0.14, 95% HPD [-0.19, -0.10]) and at 5 body lengths (ΔELPD = -0.50, 95% HPD [-0.59, -0.40]).

### Influence of subgroup composition on approaches

Leave-one-out model comparison showed that the full model with interaction and the one without had nearly identical predictive performance (ΔELPD = -0.1, SE = 1.5), whereas the null model fit substantially worse than the full model (ΔELPD = -44.9, SE = 9.6). This indicates that subgroup size and infant presence are important predictors of approach rates, but the interaction between them did not meaningfully improve model fit. In the model without interaction, approach probability increased with subgroup size (β = 0.34, 95% CI [0.18, 0.50]) and was higher when an infant was present (β = 0.69, 95% CI [0.40, 0.98]), showing clear positive main effects of both variables (**Fig.4**).

**Figure 4.**
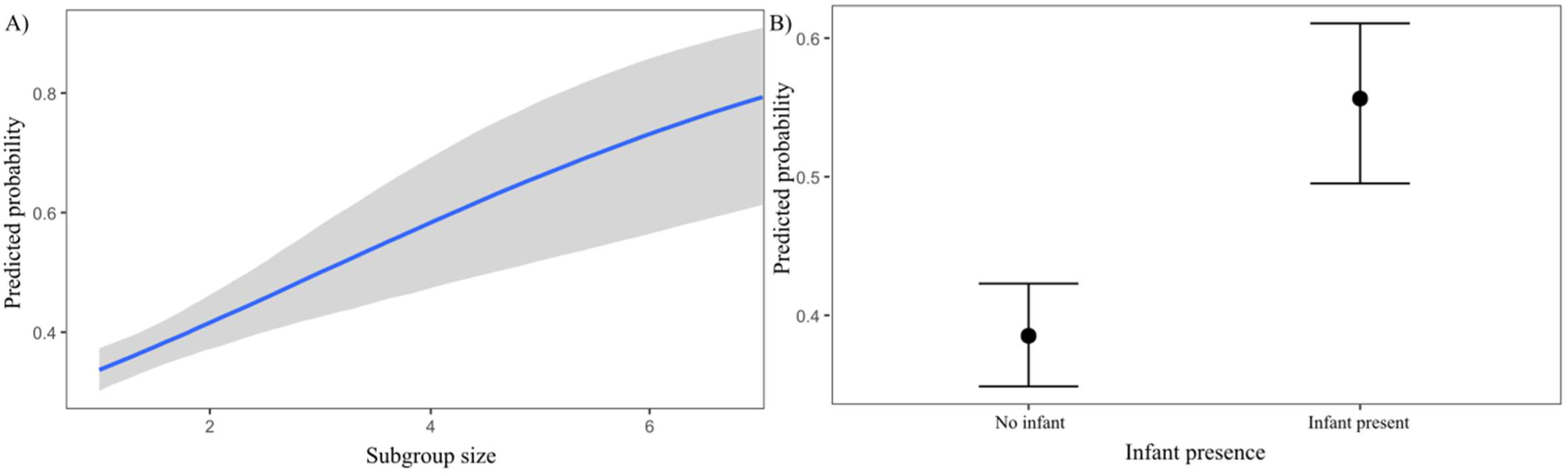
Conditional effects of subgroup size and infant presence on approach probability in Macaca nigra at Tangkoko, Indonesia (September 2025 - May 2026) **(A)** Conditional effect of subgroup size on the predicted probability of being approached, with the grey band representing the 95% credible interval around the fitted values. **(B)** Conditional effect of infant presence on the predicted probability of being approached. Points show the model-based predicted probabilities, and vertical bars represent the 95% credible intervals.

### Adult-only

We then extracted the approaches in which no infant or juvenile was present or approached, resulting in 214 adult-only approaches over 1592 approaches (∼ 13,4%). Approach events were mostly concentrated in single-individual subgroups, which were approached 191 times compared with 26 times not approached. (**Fig.5**). Approaches involving subgroups of two individuals were less frequent, with 22 approached events and only 1 non-approached event, while subgroups with 3 were rare overall, with only one event in each category. Adults approached other adults essentially when they are isolated.

**Figure 5.**
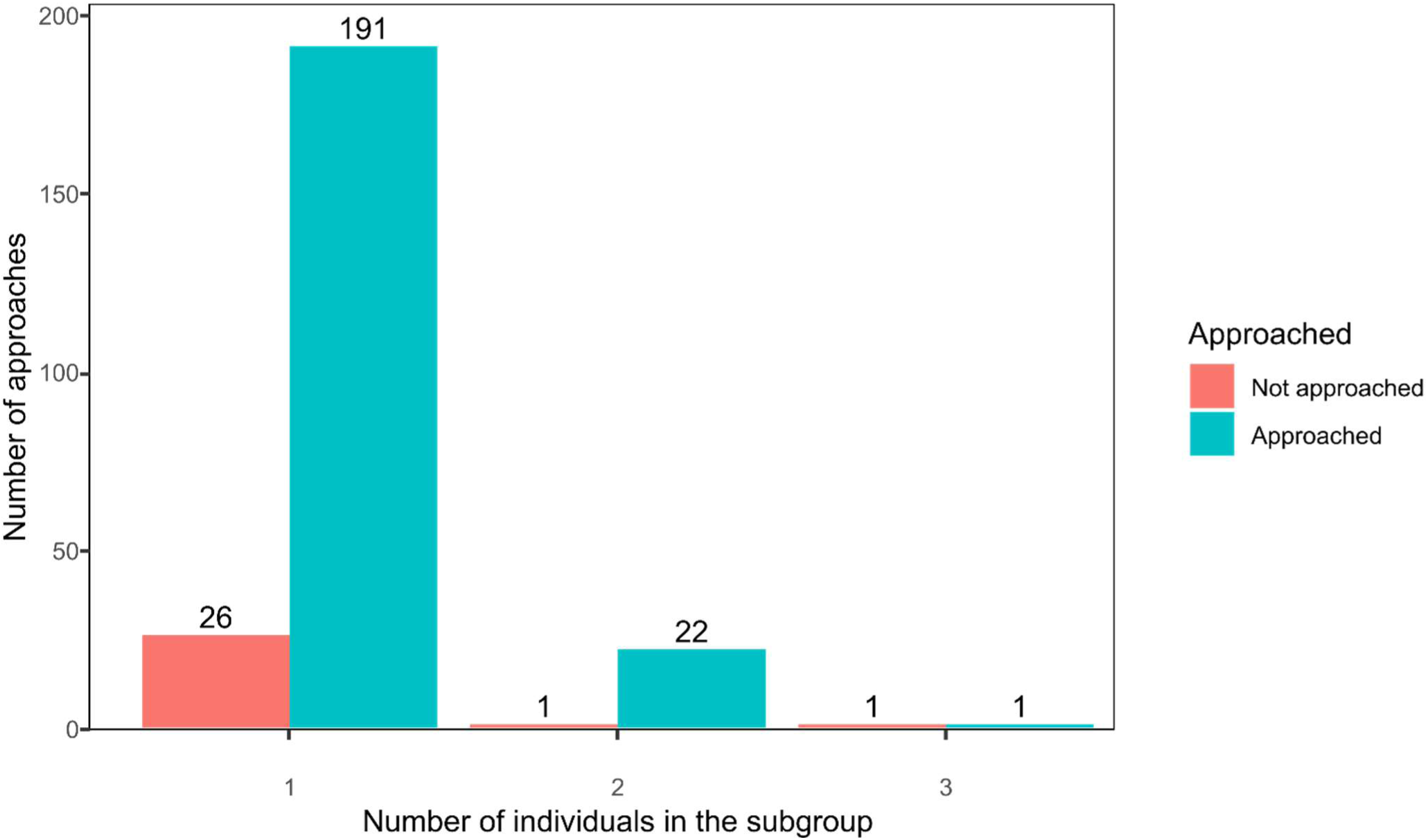
Approach occurrence by subgroup size in adult-only approaches in *Macaca nigra* at Tangkoko, Indonesia (September 2025 - May 2026) Number of approaches and non-approaches across subgroup sizes among adult-only subgroups. Bars show the number of observations for each subgroup size available during an approach, separated by approach outcome: not approached and approached.

### Subgroup choice when infants are in the pool

Finally, we extracted approaches when an infant was present in at least one subgroup in the audience (322 events), to assess whether this subgroup was more or less approached relatively to subgroups without infants. Subgroups with infants were preferentially approached (75% vs 25%; exact binomial test: p < 0.001; 95% Clopper-Pearson Credible Intervals [70%, 80%]; **Fig.6**), demonstrating strong, non-random infant-directed social selectivity during approaches.

**Figure 6.**
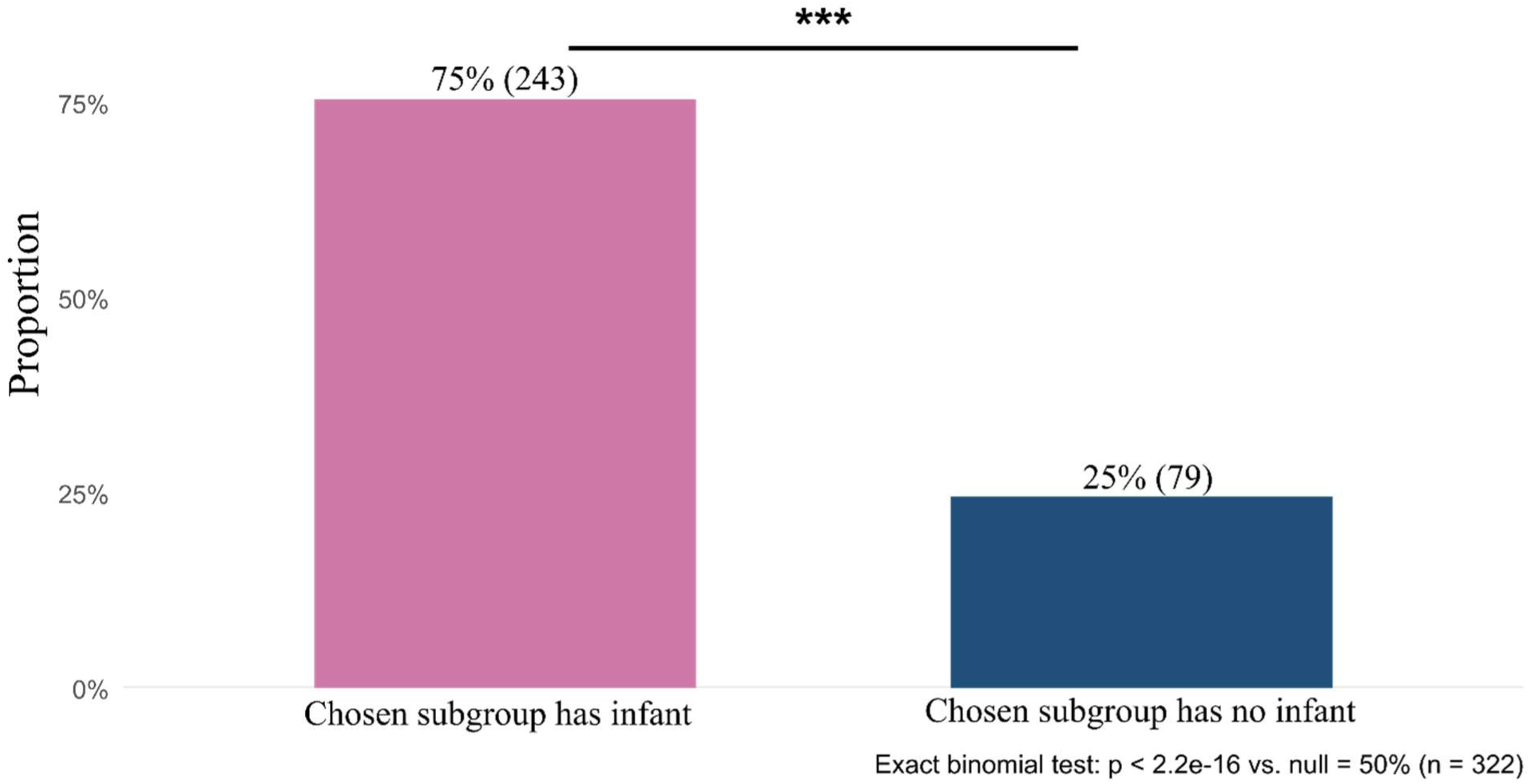
Proportion of approach directed towards groups with and without infants (September 2025 - May 2026). Bar plot of the proportion of approach choices directed toward subgroups with versus without infants.

## Discussion

Overall, our results suggest that approach decisions in crested macaques are not random but are modulated by the social context, integrating behavioural cues, social density, and infant presence. The social setting may invite, encourage or discourage approaches, depending on its local characteristics. Macaques appear to perceive neutral, affiliative, and grooming contexts as opportunities for approach, as these behaviours were more frequently associated with approach events. Individuals were also more likely to approach conspecifics that were feeding or engaged in affiliative behaviours. Although approaches most often targeted solitary individuals or dyads, when several subgroups were available, individuals preferentially approached the densest subgroups, particularly those containing an infant.

### Behavioural cues as indicators of partner availability

Behavioural cues seem to shape approach decisions by signalling the immediate availability of potential partners, conveying who may be willing to interact. At the contextual level, approach events were associated with distinct behavioural states compared to baseline activity, characterized by higher frequencies of neutral, affiliative behaviours, and grooming, and lower levels of foraging and self-grooming. These patterns suggest that approaches were embedded in socially permissive situations, rather than occurring randomly within ongoing activities. In this sense, behavioural context may constrain partners’ choice, shaping when interactions are likely to be initiated.

Within these contexts, however, partner choice appears to depend on more specific behavioural signals. Approached individuals were more likely than bystanders to display affiliative behaviours, particularly lipsmacking and affiliative facial expressions directed towards the approacher, which are known to increase the likelihood of subsequent affiliative contact (Micheletta et al., 2013). Such signals likely act as invitations to interact, reducing uncertainty and facilitating positive interaction. Similarly, partners were more often engaged in feeding behaviour, which may attract approaches because of the potential for food-related information gathering, even if it can increase competition. Macaques, like other primate species, are known to smell the mouth of an individual eating (i.e., mouth sniffing), usually to get information on the food location and quality (*Macaca tonkeana*: Drapier, Chauvin & Thierry, 2002; Chauvin & Thierry, 2005; *Mandrillus sphinx*, *Mandrillus leucophaeus*, and *Papio anubis*: Laidre, 2009; *Chlorocebus pygerythrus*: Dongre et al., 2024). Together, these findings indicate that macaques preferentially target individuals whose current activity provides clear social or ecological incentives.

In contrast, individuals already engaged in grooming were less likely to be approached, as they were more frequently found in the audience than among selected partners. This suggests that ongoing grooming may reduce accessibility, with macaques avoiding the potential costs of interrupting established interactions. By interrupting other’s grooming, approachers may risk direct aggression from the two grooming partners, and weaken their relationship with the partners involved, as well as subsequent grooming opportunities by interrupting others’ grooming. Unlike in chimpanzees and sooty mangabeys, where individuals may actively intervene in grooming interactions and sometimes disrupt ongoing bouts (Mielke et al., 2017), these results indicate a tendency to favour partners who are not already socially occupied, reinforcing the idea that interactional availability is dynamically evaluated. From this perspective, avoiding individuals who are currently engaged in grooming activities can be seen as a precautionary strategy to minimise social risks, as shown during socio-economics dilemmas in Tonkean macaques (Ballesta & Meunier, 2023). Future work could investigate what happens after approaching grooming individuals, for example whether such approaches escalate, lead to successful integration into the grooming bout, or result in displacement, to better understand how macaques balance these costs and opportunities in real time. Because grooming interference occurs in other macaque species of different social tolerance levels (Mielke et al., 2021; Thierry et al., 1990), such studies could also clarify whether crested macaques genuinely differ in their propensity to interfere in grooming bouts, or whether it was rare in our dataset, or pattern-specific to this group.

Importantly, neutral behaviours such as resting or locomotion alone had no clear effect on the variables of interest, suggesting either that availability is not fully captured by these categories or that individuals do not perceive these behaviours as cues of social availability or unavailability. Instead, at these neutral states, macaques could rely on more nuanced cues, such as body orientation, subtle postural signals, or recent interaction history to assess whether an individual is approachable. Taken together, these results suggest that a combination of behavioural context and interactional cues guides partner choice. Rather than approaching any conspecific, macaques seem to favour partners whose current behaviour indicates that they may be accessible and receptive to interaction. In this sense, individuals appear to perceive social affordances, adjusting their approach decisions to the immediate behavioural situation of potential partners.

### Rare but meaningful contexts: aggression and sex

Aggressive and sexual behaviours represent specific but infrequent contexts in which approach decisions may take on a different functional meaning. Model predictions suggested that approached individuals were more likely to display these behaviours, potentially reflecting situations in which mating opportunities or ongoing conflicts drive approaches. In the case of aggression, several non-exclusive mechanisms may explain this pattern: individuals may approach to initiate aggression, to seek protection or to provide coalitionary support. Coalitionary support is defined as “a focal individual intervening aggressively or peacefully in support of another individual or receiving such an intervention itself during an aggressive interaction with another individual” (Duboscq et al., 2014, 2017; Petit & Thierry, 1994). In such cases, the elevated rates of aggression in partners likely reflect the interactional context rather than a general aggressiveness in approaching individuals. Approached individuals never displayed sexual behaviours during our observations: behaviours that invited sexual interaction, such as sexual presentation, or the “sexual jaw-movement”, named jaw wobbling, displayed by males to solicit mating (Clark et al., 2022; Thierry et al., 2000), were classified in affiliative behaviours. No individuals engaged in mating were approached during our data collection. These two behavioural states were extremely rare in our dataset, representing only a very small fraction of observed approaches, and no sexual behaviours were directly recorded during approach events. As a result, model estimates for these categories should be interpreted with caution, as they are likely sensitive to low sample size and prior specification. Aggressive and sexual contexts frequently involve specialised interactions that seem to diverge from the main patterns we describe; targeted studies would clarify whether these contexts follow separate decision rules. Altogether, our results indicate that macaques can flexibly adjust their approach behaviour in high-stakes social contexts such as conflict or mating, but these situations did not appear to play a major role in shaping approach dynamics in our dataset.

### Social density, composition and spatial influences on partner choice

Social density, composition and spatial distribution appear to shape approach decisions, influencing not only approach occurrences but also which individuals are ultimately targeted. At a broad scale, approaches took place in dense social contexts, as the number of neighbours was consistently higher during approach events than during focal observations across all distance categories. In primates, group size reflects a balance between social opportunities and constraints: while subgroups with more conspecifics can increase opportunities for positive interactions, they may also intensify competition, reduce access to specific partners, and increase possible interaction disruption (Chapman & Teichroeb, 2012). Beyond competition for access to partners, close social proximity may also increase the likelihood that ongoing affiliative interactions are interrupted by third parties (Mondragón-Ceballos, 2001). Our findings suggest that, in this group of crested macaques, density functions as an attractor, consistent with their tolerant social style and previously described dense and evenly distributed proximity or grooming networks (Balasubramaniam et al., 2018; Duboscq et al., 2013).

However, partner choice within these dense contexts was constrained by spatial distribution. When comparing approached individuals to surrounding bystanders, partners were typically more spatially isolated at greater distances (one and five body lengths), indicating that individuals preferentially targeted those who were locally isolated, likely to minimise the risk of interruption. This pattern reveals a finer scale selection process: macaques may be drawn to dense social settings, but within them they favour partners who are temporarily on their own, away from ongoing social engagement, and therefore easier to interact with. Importantly, our data only capture approach decisions, not the interactions that may or may not follow, so we interpret proximity as indicating social opportunity rather than confirmed social engagement. In this light, our results suggest that individuals use their immediate social context to navigate toward partners that offer more favourable opportunities for interactions, while still operating within crowded neighbourhoods.

An apparent exception to this pattern emerged at the closest spatial scale, where partners had more neighbours in body contact than bystanders. This effect, however, was driven by the presence of infants. Mother–infant dyads, characterised by high levels of body contact, were disproportionately approached, and removing infants from the analysis eliminated the difference between partner and audience. This supports the idea that infant presence modifies local spatial configurations by increasing the attractiveness of the subgroup they are in, in line with previous findings on infant attractiveness in primates (Dunayer & Berman, 2017; Maestripieri, 1994; Silk et al., 2013). We also note that these analyses include only approaches that occurred when more than one subgroup was present (i.e., when individuals had a choice among partners). Approaches directed at mother–infant dyads in the absence of other subgroups were also frequent, highlighting the strong infant attractiveness effect. Taken together, these results suggest that crested macaques make trade-offs between social attraction and interactional accessibility. While they preferentially enter denser social environments, they tend to select locally isolated partners within these contexts, preferring to approach isolated individuals and mother-infant dyads. This interpretation is further supported by the observation that, in adult-only contexts, individuals predominantly approached isolated partners. This pattern may indicate a genuine preference for partners who are temporarily isolated, but further work is needed to determine whether individuals actively choose isolated partners or whether this bias arises passively from the underlying spatial structure of the group.

### Integrating affordances with social relationships

In the studied crested macaques, our findings suggest that immediate social context shapes the social affordances that structure partner choice. Contextual cues such as social density, behavioural state, and infant presence appear to define a set of available partners, constraining who can be approached at a given moment. Within this constrained set, however, approach decisions are unlikely to be opportunistic and may depend on additional factors, such as social relationships, past experience and individual preferences, which remain to be integrated in our line of study.

Although the present study primarily highlights contextual cues, individual personality differences may constitute an additional layer of decision-making. Differences in traits such as sociability, connectedness, or aggressiveness (Neumann et al., 2013) may influence how individuals perceive and use social affordances, leading some to preferentially engage in interactions while in dense social settings, while others may avoid them. In this sense, personality may modulate sensitivity to context, shaping how opportunities are translated into actual interactions.

A key remaining question finally concerns how these temporary affordances interact with stable social relationships in guiding approach decisions. Future analyses incorporating rank, kinship, and social bonds will be essential to disentangle whether these factors act independently, hierarchically, or in combination with immediate affordances. For instance, strong social bonds may override constraints imposed by temporary unavailability, allowing individuals to approach partners who are otherwise engaged, whereas hierarchical distance might inhibit approaches despite apparent accessibility. Indeed, in females crested macaques, stronger bonds are formed between closely ranked but not kin or age peers, and stronger bonds were more equitable but less predictable than weaker bonds (Duboscq et al., 2017). This suggests a temporally changing social landscape which may be challenging to navigate without correct social knowledge.

In the crested macaque population we studied, our approach-based analyses allowed us to identify which configurations of behaviour and local social context are treated as opportunities for social interaction. Partner choice emerges as a flexible, context-dependent process, where immediate social affordances delimit the set of potential interaction partners, while factors such as social relationships or individual personality likely shape which of these opportunities is actually taken

## Supporting information

Electronic Supplementary Materials

## Acknowledgments

We thank the members of the Macaca Nigra Project and the assistants for their help with data collection. We gratefully acknowledge the Department for the Conservation of Natural Resources (BKSDA), and the National Research and Innovation Agency (BRIN) for permitting us to carry out this research. We are grateful to Askhari Dg. Masikki S.Hut, head of the BKSDA Sulut for providing local administrative and logistic support without which this research would not have been possible. We finally thank Mathilde Haeuser for providing the macaque illustration in Figure 1.

## Data accessibility

The data will be available on the InDoRES platform (https://data.indores.fr), a secure institutional and thematic data repository.

## Authors’ contribution

A.P. and F.B. designed the study and data collection protocol, with the supervision of J.D., C.G., and S.B.; A.P. and M.J.B collected the behavioural data; A.P., with the supervision of J.D., C.G., and S.B, designed and conducted the data analysis and drafted the manuscript; A.P., J.D., C.G., S.B., F.B and D.P-F reviewed the manuscript and approved the submitted version.

