## Supplementary material for "A Social Affordance Framework for Understanding Approach Decisions in Wild Crested Macaques (*Macaca nigra*)": Electronic Supplementary Materials

**Electronic Supplementary Material**

| **Behaviour** | **Difference (Focal - Approach)** | **95% HPD Lower** | **95% HPD Upper** | **Significance** |
| --- | --- | --- | --- | --- |
| Neutral | -5.507376 | -7.0527 | -3.8812 | **Yes** |
| Autogrooming | 1.069017 | 0.4144 | 1.6750 | **Yes** |
| Affiliative | -1.069354 | -1.6823 | -0.4397 | **Yes** |
| Aggressive | 0.086630 | -0.1159 | 0.2871 | No |
| Feeding | 0.513270 | -1.5279 | 2.5138 | No |
| Foraging | 4.691755 | 2.1285 | 7.3382 | **Yes** |
| Grooming | -3.764165 | -5.7318 | -2.1058 | **Yes** |
| Sexual | 0.000471 | -0.0878 | 0.0707 | No |

**Table 1. Pairwise contrasts of estimated marginal means for predicted behaviour rates across contexts (Model: Focal vs. Approach) and across roles (Model: Partner vs. Audience) in *Macaca nigra* at Tangkoko, Indonesia (September 2025 - May 2026).** Differences (Focal versus Approach) with 95% highest posterior density intervals. Effects were considered significant when the interval did not include zero. The highest posterior density intervals were obtained from posterior samples from the Bayesian models, and represent the shortest intervals containing 95% of the posterior probability mass. Positive values indicate higher rates in the focal context than in the approach context.

| **Behaviour** | **Difference (Partner - Audience)** | **95% HPD Lower** | **95% HPD Upper** | **Significance** |
| --- | --- | --- | --- | --- |
| Neutral | -0.3732 | -0.8134 | 0.0699 | No |
| Autogrooming | 0.0898 | -0.1004 | 0.2755 | No |
| Affiliative | 0.4731 | 0.2401 | 0.6985 | Yes |
| Aggressive | 0.1084 | 0.0214 | 0.2105 | Yes |
| Feeding | 0.8644 | 0.3032 | 1.4370 | Yes |
| Foraging | -0.4494 | -1.1638 | 0.3134 | No |
| Grooming | -0.9652 | -1.6200 | -0.3267 | Yes |
| Sexual | 0.0866 | 0.0118 | 0.1683 | Yes |

**Table 2. Pairwise contrasts of estimated marginal means for across roles (Partner vs. Audience) in *Macaca nigra* at Tangkoko, Indonesia (September 2025 - May 2026).**

Differences (Partner versus Audience) with 95% highest posterior density intervals. Effects were considered significant when the interval did not include zero. The highest posterior density intervals were obtained from posterior samples from the Bayesian models, and represent the shortest intervals containing 95% of the posterior probability mass. Positive values indicate higher rates in the partner than in the audience.

| **Distance category** | **Difference (Focal - Approach)** | **95% HPD lower** | **95% HPD upper** | **Significance** |
| --- | --- | --- | --- | --- |
| Body contact | -3.19 | -3.96 | -2.38 | **Yes** |
| 1 body length | -1.68 | -2.24 | -1.14 | **Yes** |
| 5 body lengths | -3.65 | -4.91 | -2.37 | **Yes** |

**Table 3. Pairwise contrasts of estimated marginal means for distance category across contexts (Model: Focal versus Approach) in *Macaca nigra* at Tangkoko, Indonesia (September 2025 - May 2026).** Differences with 95% highest posterior density intervals. Effects were considered significant when the interval did not include zero. The highest posterior density intervals were obtained from posterior samples from the Bayesian models, and represent the shortest intervals containing 95% of the posterior probability mass. Positive values indicate higher rates in the focal context than in the approach context.

| **Distance category** | **Difference (Partner − Audience)** | **95% HPD lower** | **95% HPD upper** | **Significance** |
| --- | --- | --- | --- | --- |
| Body contact | 0.113 | 0.0627 | 0.164 | **Yes** |
| 1 body length | -0.144 | -0.1878 | -0.101 | **Yes** |
| 5 body lengths | -0.496 | -0.5853 | -0.398 | **Yes** |

**Table 4. Pairwise contrasts of estimated marginal means for distance category across roles (Model: Partner versus Audience) in *Macaca nigra* at Tangkoko, Indonesia (September 2025 - May 2026).** Differences with 95% highest posterior density intervals. Effects were considered significant when the interval did not include zero. The highest posterior density intervals were obtained from posterior samples from the Bayesian models, and represent the shortest intervals containing 95% of the posterior probability mass. Positive values indicate higher rates in the focal context than in the approach context.

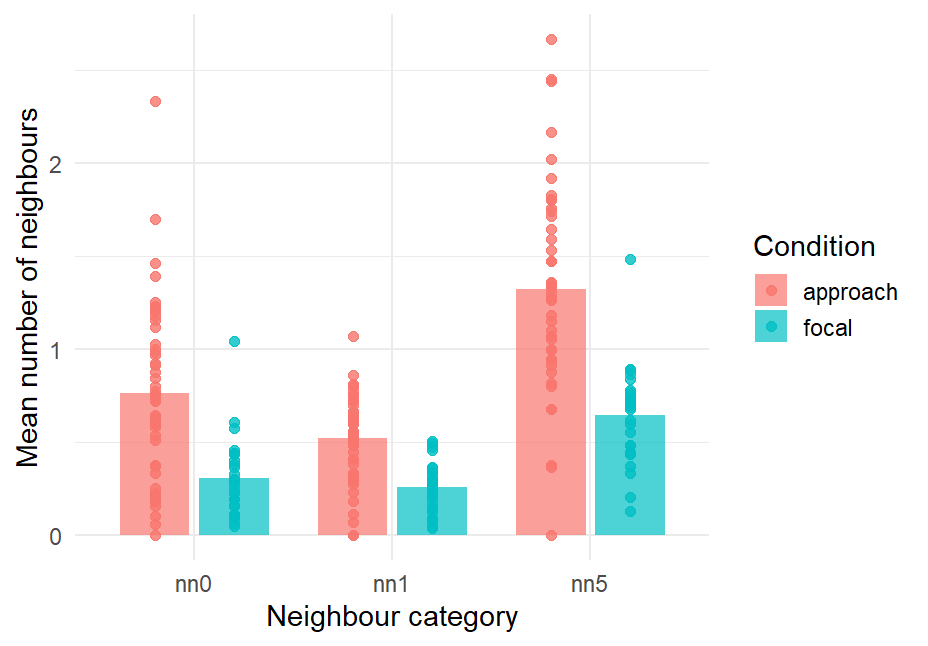

**Figure 1. Mean number of neighbours by condition and neighbour category in *Macaca nigra* at Tangkoko, Indonesia (September 2025 - May 2026).** Comparison of focal and approach observations across body contact (nn0), 1 body length (nn1), and 5 body lengths (nn5), with dots showing individual means.

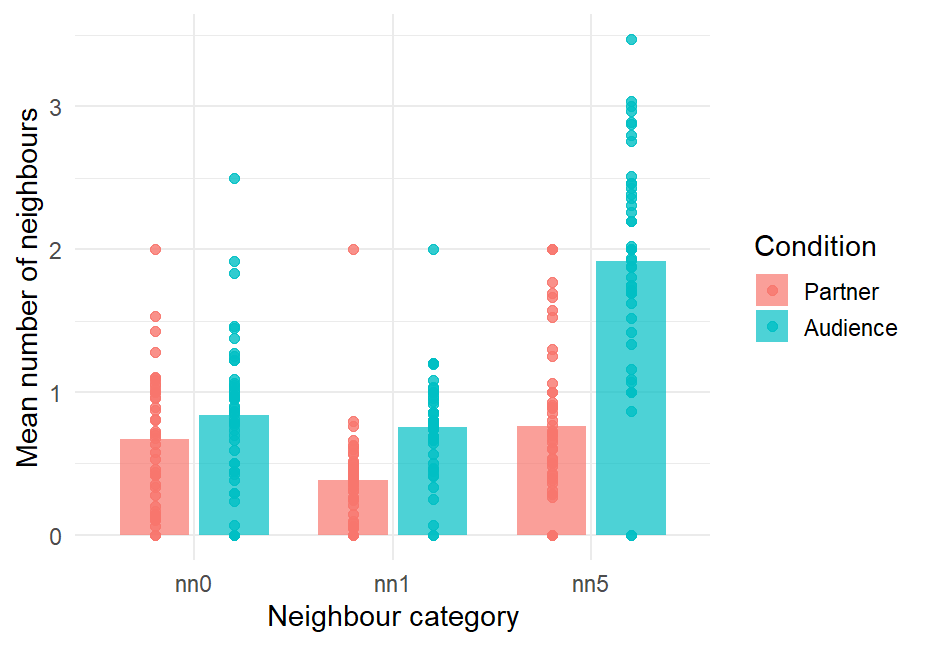

Role

Role

**Figure 2. Mean number of neighbours by condition and neighbour category in *Macaca nigra* at Tangkoko, Indonesia (September 2025 - May 2026).** Comparison of partner and audience observations across body contact (nn0), 1 body length (nn1), and 5 body lengths (nn5), with dots showing individual means.
